# Improving angiogenesis ameliorates the efficacy of ASO-based exon-skipping for the treatment of Duchenne muscular dystrophy

**DOI:** 10.1101/2025.08.21.670323

**Authors:** Mathilde Blitek, Cécile Gastaldi, Mathilde Doisy, Olivier Le Coz, Marion David, Xaysongkhame Phongsavanh, Sameh Ben Aicha, Luis Garcia, Alessio Rotini, Gilles Pagès, Aurélie Goyenvalle

## Abstract

Duchenne muscular dystrophy (DMD) is a severe X-linked disease caused by mutations in the *DMD* gene, resulting in the absence of functional dystrophin. Antisense oligonucleotide (ASO)-based therapies aim to restore the open reading frame and produce a truncated but functional dystrophin protein. Although several ASOs are FDA-approved, dystrophin restoration in patient biopsies remains low, underlining the need to improve ASO efficacy. One major limitation is poor ASO biodistribution to skeletal muscle, influenced by both ASO chemistry and pathological features of dystrophic tissue. In DMD patients and *mdx* mice, microvascular abnormalities and impaired angiogenesis likely restrict ASO delivery. Here, we hypothesized that enhancing muscle vascularization could improve ASO biodistribution and therapeutic outcomes. *Mdx* mice were treated with a pro-angiogenic treatment prior to ASO administration targeting exon 23 of dystrophin pre-mRNA. Angiogenic stimulation increased capillary density and improved ASO delivery, exon skipping, and dystrophin expression compared to ASO alone. These molecular improvements were associated with increased myofiber size, larger mean cross-sectional area, and decreased serum myomesin levels, without signs of toxicity. This study provides proof-of-concept that promoting angiogenesis can enhance the efficacy of ASO-based treatments, offering a complementary strategy to improve therapeutic outcomes in DMD.

## INTRODUCTION

Duchenne muscular dystrophy (DMD) is a severe X-linked neuromuscular disorder characterized by progressive muscle degeneration. It affects approximately one in 5,000 male newborns and leads to a rapid loss of ambulation, followed by cardiac and respiratory complications that ultimately result in premature death.^1^ DMD is caused by mutations in the *DMD* gene, with deletions accounting for approximately 65% of cases.^2^ These mutations disrupt the reading frame, leading to the introduction of a premature stop codon and the subsequent loss of dystrophin expression.^3^ Dystrophin is a key membrane-associated protein that connects the cytoskeleton to the extracellular matrix, playing a crucial role in protecting muscle fibers from the mechanical stress associated with contractions. In the absence of functional dystrophin, the sarcolemma becomes fragile and prone to rupture during muscle contractions.^4^

Among the therapeutic strategies under development, exon skipping has emerged as a promising approach for restoring dystrophin expression. Since most DMD-causing mutations result in frameshifts, selectively excluding specific exons can help re-establish an open reading frame and produce a truncated yet functional dystrophin protein.^5^ This technique relies on antisense oligonucleotides (ASOs), which are short synthetic nucleic acid sequences designed to bind to pre-mRNA targets and modulate splicing. Over the years, ASO chemistries have evolved to improve stability and efficacy, replacing unmodified DNA or RNA sequences.^6^ Early clinical trials investigated 2′O-methyl (2′O-Me) and phosphorodiamidate morpholino oligomers (PMOs), with some PMO-based therapies being approved in the United States and Japan for skipping exons 45, 51, and 53.^7–10^ However, these ASOs face challenges such as limited distribution to muscle tissues and relatively low levels of dystrophin restoration.^11^ To overcome these limitations, next-generation ASOs are being developed. Among them, tricyclo-DNA (tcDNA) oligonucleotides, particularly when conjugated to palmitic acid, have demonstrated strong therapeutic potential in the *mdx* mouse model.^12–14^ Their improved RNA affinity and enhanced biodistribution make them promising candidates for DMD treatment.

Nevertheless, given the complexity of DMD, meaningful functional improvements will likely require combinatorial strategies, regardless of the therapeutic modality employed. In the context of muscular dystrophies, modulating disease pathophysiology itself may play a crucial role. Improving muscle histopathology, even without changes in dystrophin levels, can lead to enhanced muscle function.^15^ In this study, we focused on the microvasculature which is impaired in both DMD patients and *mdx* mouse models.

Angiogenesis is a highly orchestrated physiological process regulated by a fine-tuned equilibrium between pro-angiogenic and anti-angiogenic cues, through which neovasculature arises from pre-existing blood vessels. This process is indispensable for sustaining adequate oxygenation and metabolic exchange, both of which are critical for optimal skeletal muscle function and systemic homeostasis. In DMD, both patients and *mdx* mice exhibit pronounced microvascular pathology, characterized by diminished capillary density, endothelial cell swelling and pallor, aberrant vascular architecture, and frequent capillary occlusions.^16–18^ These abnormalities collectively exacerbate tissue hypoxia and promote focal myofiber necrosis. Moreover and more importantly in the context of therapeutic interventions, compromised microvasculature may significantly compromise the biodistribution and tissue penetrance of ASOs, thereby limiting the therapeutic efficacy of ASO-based intervention.

Among known pro-angiogenic factors, vascular endothelial growth factor A (VEGFA) is one of the most potent.^19^ Through its interaction with the VEGF receptor 2 (VEGFR2), VEGFA activates key signaling pathways that drive endothelial cell proliferation, migration, survival, and vascular permeability, leading to the formation of new capillary networks.^20^ In skeletal muscle, VEGFA enhances vascularization, reduces local inflammation, and improves muscle regeneration, features that could be beneficial in the context of DMD.^21,22^ However, local doses of recombinant VEGFA have yielded modest benefits and raised safety concerns, including inflammatory responses and disorganized or excessive angiogenesis.^23,24^ These outcomes highlight the complexity of establishing a therapeutic window for VEGFA that maximizes angiogenic benefit while minimizing potential deleterious consequences.

In the present study, we investigated a recently characterized VEGFA isoform, termed VEGFA new form (VEGF-NF) or Lymphatic and Vascular Resistant Factor (LVRF), a nomenclature that reflects its newly identified functional properties. This isoform arises from the activation of an alternative upstream splice acceptor site within intron 7 of the *VEGFA* gene.^25^ This splicing event introduces a 23-base pair intronic sequence, referred to as the NF domain, and gives rise to a novel open reading frame that extends into a region typically considered part of the 3′ untranslated region (3′UTR). This isoform promotes the formation of both functional blood and lymphatic vessels, enabling a more controlled and physiologically balanced angiogenic response with an improved safety profile..^25^

We hypothesized that improving the vascular microenvironment in DMD could not only ameliorate vascular deficiencies and improve muscle perfusion and oxygenation, but also facilitate the biodistribution and uptake of ASOs, ultimately increasing their therapeutic potential. To this aim, we vectorized the human (hLVRF) and murine LVRF (mLVRF) isoforms, under the muscle specific promoter CK8, into an AAV9 (adeno-associated virus of serotype 9) vector enabling particularly efficient gene transfer into muscles. We then evaluated the therapeutic impact of combined LVRF gene transfer with tcDNA-ASOs targeting exon 23 of the *Dmd* gene in *mdx* mice.

## RESULTS

### Effect of hLVRF on angiogenesis after intramuscular injection in *mdx* mice

We first assessed the therapeutic potential of hLVRF on angiogenesis in a preliminary study after intramuscular delivery. Adult *mdx* mice received a single intramuscular injection of 5×10¹¹ viral genomes per tibialis anterior (TA) of AAV9 encoding the previously described hLVRF isoform^25^ (**Figure S1A**). Four weeks following AAV-mediated LVRF pre-treatment, mice received intravenous administration of ASO therapy (50 mg/kg/week), delivered once weekly for four consecutive weeks to promote dystrophin restoration (**Figure 1A**). TA muscles were harvested one week after the end of ASO treatment for analysis. To evaluate the impact of hLVRF expression on blood vessels, we first performed CD31 immunostaining on TA cross-sections. In PBS-treated *mdx* mice, a high proportion of muscle fibers were associated with fewer than four capillaries, and a lower proportion with more than five vessels, compared to wild-type (WT) animals (**Figure 1B**). In contrast, hLVRF-treated mice displayed an improved vascular phenotype. Quantitative analysis of vascularization, measured as the mean number of vessels per muscle fiber, revealed a significant reduction in PBS-treated *mdx* mice compared to WT controls. Notably, hLVRF administration significantly increased vascular density relative to PBS-treated *mdx* mice, restoring vessel numbers to levels that were not statistically different from WT (**Figure 1C**). Given the previously reported ability of LVRF to uniquely promote the formation of both blood and lymphatic vessel, we further investigated its lymphangiogenic potential *in vitro* using a 3D lymphatic endothelial cell (LEC) sprouting assay. Recombinant human LVRF, VEGFA, or VEGFC (200 ng/mL) were tested as stimuli. Sprouting was quantified by counting sprout-positive beads per condition (Fig S1B). While VEGFA failed to induce LEC sprouting, both VEGFC and LVRF significantly promoted sprout formation compared to control and VEGFA-treated conditions (**Figure S1C**). This lymphangiogenic activity was also confirmed *in vivo*, as *Lyve1* (Lymphatic vessel endothelial receptor 1) mRNA levels were significantly increased in TA muscles injected with hLVRF compared to PBS-treated controls (**Figure S1D**).

**Figure 1:**
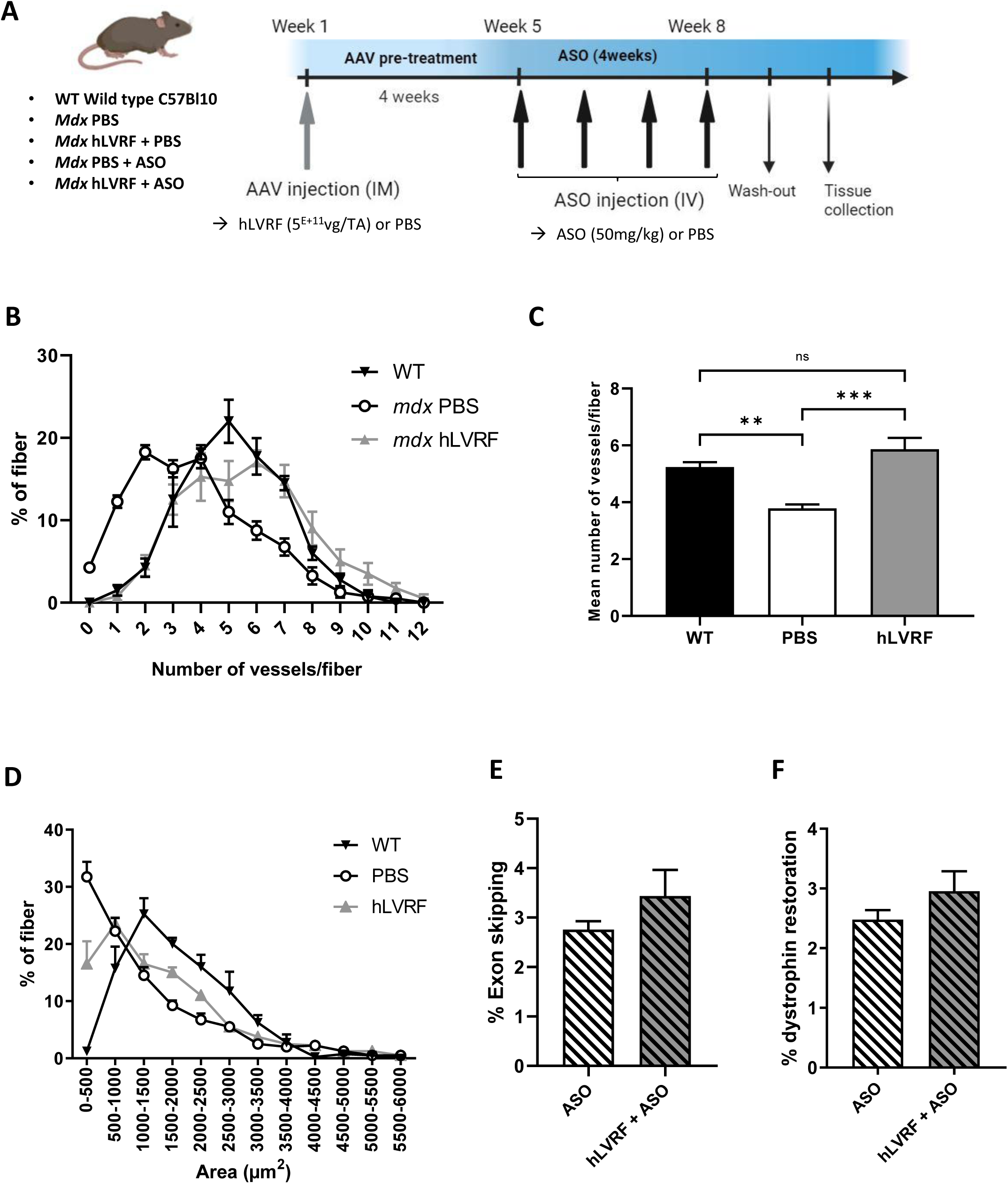
Effect of human LVRF expression on angiogenesis. **(A)** Timeline of the injection protocol. Groups of male *mdx* mice received a single intramuscular injection of AAV9 encoding the hLVRF isoform (5E+11 vg/muscle) or PBS in the tibialis anterior (TA) muscle. Four weeks after this pre-treatment, mice were injected intravenously with ASO (50 mg/kg/week) or PBS during 4 weeks. Age-matched C57BL/10 wild-type mice were used as controls (WT). One week after the last injection, tissues were collected as described in the material and method. **(B)** Distribution of the proportion of fibers as a function of the number of associated blood vessels. **(C)** Quantification of the mean number of blood vessels per myofiber. (**p<0.01; ***p<0.001, analyzed by one-way ANOVA). **(D)** Representation of the percentage of fibers as a function of area including several intervals. The data regarding angiogenesis and fiber size were obtained through CD31 and laminin co-staining of tibialis anterior (TA) cross-sections. **(E)** Quantification by qPCR of exon 23 skipping in TA muscles. **(F)** Quantification of dystrophin expression by western-blot (WB) on TA lysates. Results are expressed as the mean ± SEM (n=4 per group).

Besides the improvement of the microvascular network, we also assessed whether hLVRF expression could influence muscle fiber size, another hallmark of the dystrophic phenotype. TA muscles from *mdx* mice indeed exhibit a higher number of small fibers (<500 µm^2^) and a reduced number of large fibers compared to WT mice, indicative of strong regeneration. The muscle fiber size distribution in hLVRF-treated muscles more closely resembled that of WT controls, characterized by a 15% reduction in the proportion of small fibers (<500 µm²) and a 6% increase in fibers with cross-sectional areas between 1500 and 2000 µm² (**Figure 1D**).

Given the observed improvement in microvascular architecture, we hypothesized that ASO-based therapy would exhibit enhanced efficacy in restoring dystrophin expression, owing to improved biodistribution and delivery to target muscle tissues. Quantification of exon skipping revealed a 25% increase in TA muscle of mice treated with hLVRF and ASO compared to ASO treatment alone (**Figure 1E**). This enhancement in exon skipping translated into 20% increase in dystrophin restoration in hLVRF+ASO-treated muscles (**Figure 1F**). Although these changes did not reach statistical significance, the results suggest that LVRF has the capacity to enhance angiogenesis in TA muscles, thereby facilitating improved dystrophin restoration via exon skipping and contributing to amelioration of muscle histopathology in *mdx* mice.

### Systemic Delivery and Comparative Evaluation of mLVRF and VEGFA₁₆₄ in *mdx* Mice

Encouraged by the beneficial effects observed following local delivery, we next aimed to evaluate the systemic therapeutic potential of LVRF and compare its efficacy to that of VEGFA. To this end, we cloned the murine canonical VEGFA₁₆₄ isoform (the most abundantly expressed VEGFA variant in mice) as control and the corresponding murine LVRF isoform (mLVRF), into an AAV9 under the control of the muscle specific promoter CK8. To determine the optimal systemic dose for muscle vasculature improvement, a dose–response study (1E+12 vg/kg to 3E+13 vg/kg) was then conducted using intravenous AAV injections. Despite similar copy number of viral genomes found in muscles after injection of both vectors (**Figure S2A**), RT-qPCR revealed comparable expression levels of the transgenes at 1E+13 vg/kg for VEGFA₁₆₄ and 3E+13 vg/kg for mLVRF (**Figure S2B**). At these doses, both isoforms significantly increased CD31 expression in muscles (**Figure S2C**). In addition, flow cytometry analysis on freshly isolated muscles revealed that both treatments restored endothelial cell numbers to WT levels, with a more pronounced effect observed in the mLVRF group (**Figure S2D**). We subsequently assessed myofiber distribution as a function of their associated vascularization, quantifying the number of blood vessels per fiber. Both treatments shifted the capillary distribution curve toward WT levels; however, muscles treated with mLVRF most closely approximated the WT profile, exhibiting a peak of 22% of myofibers associated with five blood vessels (**Figure S2E**). Analysis of muscle fiber size distribution profiles, commonly altered in *mdx* mice, revealed that VEGFA₁₆₄-treated muscles displayed a worsened profile compared to PBS controls, characterized by an increased proportion of small fibers (<1000 µm²) and a reduced number of large fibers (>2500 µm²). In contrast, mLVRF-treated muscles displayed a distribution pattern more closely resembling that of WT, with fewer small fibers and more large fibers (**Figure S2F**). Consistent with these morphological improvements, only mLVRF treatment resulted in a significant 15% reduction in the proportion of centrally nucleated fibers, a hallmark of ongoing muscle regeneration and dystrophic pathology (**Figure S2G**). Collectively, these findings indicate that systemic delivery of mLVRF confers histopathological benefits even as a monotherapy and underscore its therapeutic potential, particularly in the context of combinatorial strategies involving ASO treatment.

Based on the dose-finding study, a dose of 3E+13 vg/kg was selected for AAV9-mLVRF in subsequent experiments. In contrast, systemic administration of AAV-VEGFA₁₆₄ was discontinued due to severe adverse effects, including mortality in several animals, even at the lower dose of 1E+13 vg/kg.

### Impact of the combined LVRF+ASO treatment on muscle microvasculature

We next explored the potential of a combined strategy using mLVRF and ASOs. Adult *mdx* mice received an intravenous injection of AAV9 encoding mLVRF at a dose of 3E+13vg/kg or a control AAV9 (scramble), followed by weekly ASO injections for 12 weeks, starting one month after AAV administration (**Figure 2A**). Given our hypothesis that enhanced vascularization may improve ASO distribution, we first evaluated the effects of mLVRF expression on microvascular architecture. To this end, we confirmed LVRF expression at the mRNA level using qPCR (**Figure 2B**). These result indicate that mLVRF expression is selectively increased in muscles injected with AAV-mLVRF. Transgene expression was also confirmed at the protein level by ELISA and showed similar up-regulation in mLVRF injected muscles (**Figure 2C**). Moreover, mLVRF overexpression correlated with increased CD31 expression (a widely used marker of vascular endothelial cells) effectively compensating for the reduced levels observed in scramble-treated *mdx* mice (**Figure 2D**). To determine whether this translated into an actual increase in vascular density, we analyzed the capillary-to-fiber ratio. CD31 immunostaining was performed on TA cross-sections to quantify the number of blood vessels associated with each myofiber (**Figures 2E-F**). As previously reported, the *mdx* scramble group exhibited a reduced number of vessels per fiber compared to WT mice. ASO treatment alone had no impact on this parameter, while mLVRF administration significantly increased the capillary-to-fiber ratio (**Figure 2F**). This was further confirmed by the mean number of vessels per fiber: the scramble group exhibited significantly fewer vessels than WT mice, while only mLVRF-treated groups showed a significant increase (**Figure 2G**). Specifically, the percentage of fibers lacking associated vessels significantly decreased in the mLVRF-treated group (**Figure 2H**), while fibers associated with five vessels increased by approximately 10% (**Figure 2I**). Building on the known lymphangiogenic potential of LVRF, we next investigated whether mLVRF modulates lymphatic vessels *in vivo* by quantifying *VEGFR3* expression, a lymphatic endothelial cell marker, via qPCR. A significant increase in VEGFR-3 expression was observed in the mLVRF+ASO group (**Figure 2J**), suggesting enhanced lymphatic vessel formation alongside improved blood vascularization. To further confirm the functionality of the newly formed blood vessels, we performed intravenous injections of fluorescently labeled-isolectin in some treated mice before their euthanasia and tissue collection, in order to visualize the vascular network in 3D after optical tissue clearing. Isolectin binds specifically to endothelial cells in perfused and functional blood vessels, allowing their detection under fluorescence microscopy. If the vessels are immature, leaky, or non-functional, isolectin rapidly diffuses out of the circulation, resulting in weak or no signal. Detection of the fluorescently labeled-isolectin revealed disorganized, tortuous blood vessels in PBS-injected muscles, whereas newly formed vessels in mLVRF-treated muscles were fully perfused and functionally integrated into the circulation (**Figure 3A**).

**Figure 2:**
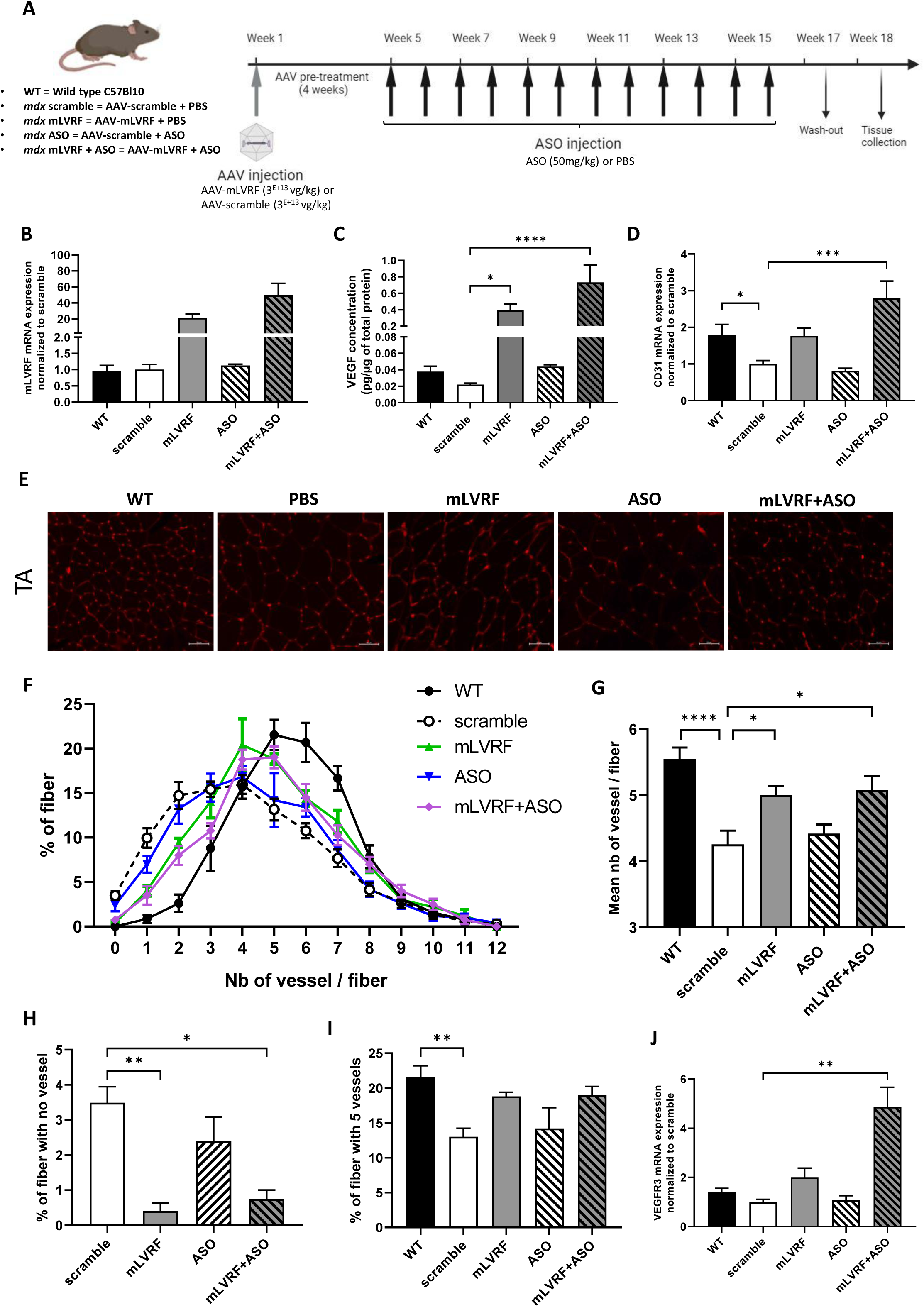
Angiogenesis improvement after mLVRF systemic injections. **(A)** Timeline of the injection protocol. Groups of 7-10 week-old male *mdx* mice were injected intravenously with a single dose AAV9 encoding the mLVRF isoform or an AAV9-scramble as control (3E+13 vg/kg) (n=4-5 mice per group). Four weeks after this pre-treatment, mice were injected intravenously with ASO (50 mg/kg/week) or PBS during 12 weeks. Age-matched C57BL/10 wild-type mice were used as controls (WT). One week after the last injection, tissues were collected as described in the material and method section. **(B)** Quantification of the expression of mLVRF performed by qPCR on gastrocnemius (GAS) total RNA. **(C)** VEGF concentration measured by ELISA on GAS protein lysates. (*p<0.05; ****p<0.0001, analyzed by one-way ANOVA). **(D)** CD31 expression quantified by qPCR on total RNA extracted from GAS muscles. (*p<0.05; ***p<0.001, analyzed by one-way ANOVA). **(E)** CD31 immunostaining of TA muscles from WT, scramble, mLVRF, ASO and mLVRF+ASO treated mice. Scale bar, 50µm. **(F)** Distribution of the proportion of fibers as a function of the number of associated blood vessels. **(G)** Quantification of the mean number of blood vessels per myofiber. (*p<0.05; ****p<0.0001, analyzed by one-way ANOVA). Proportion of fibers with no associated blood vessels **(H)** or associated with 5 vessels **(I)**. (*p<0.05; **p<0.01 analyzed by one-way ANOVA). Data regarding blood vessels were obtained from CD31 immunostaining. **(J)** Quantification of the *VEGFR3* expression by qPCR on GAS RNA. (**p<0.01, analyzed by one-way ANOVA). Results are expressed as the mean ± SEM (n=5 in WT, scramble, mLVRF and ASO groups, n=4 in mLVRF+ASO group).

**Figure 3:**
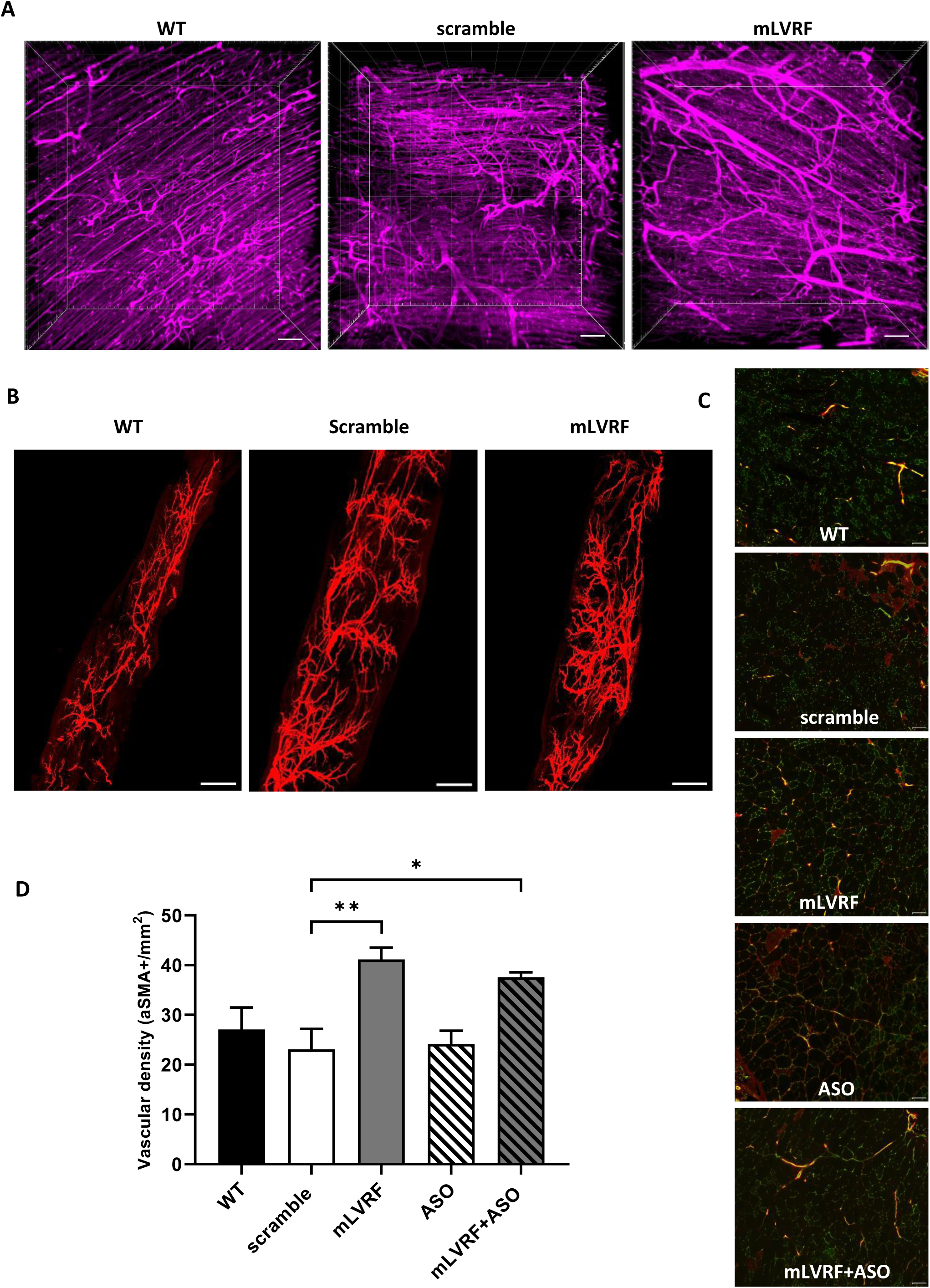
Functionality and maturation of blood vessels. **(A)** Fluorescence microscopy of TA muscle after DyLight649-isolectin intravenous injections and tissue clearing of WT, scramble and mLVRF treated mice. Scale bar, 100µm. (**B)** Fluorescence microscopy after tissue clearing and aSMA staining of EDL (extensor digitorum longus) muscle of WT, scramble and mLVRF treated mice. Scale bar, 500µm. (**C)** aSMA (red) and CD31 (green) immunostaining on transverse sections of triceps muscles from WT, scramble, mLVRF, ASO and mLVRF+ASO treated mice. Scale bar, 100µm. (**D)** Quantification of the number of aSMA+ blood vessel per mm^2^ from the aSMA and CD31 cross-section stainings (*p<0.05; **p<0.01, analyzed by one-way ANOVA). Results are expressed as the mean ± SEM (n=5 in WT, scramble, mLVRF and ASO groups, n=4 in mLVRF+ASO group).

Beyond assessing vascular functionality, we evaluated vessel maturation by staining for α-smooth muscle actin (αSMA), a marker of vascular smooth muscle cells (vSMCs), which contributes to vessel stabilization and regulation. αSMA staining combined with tissue clearing of TA muscles suggested a higher density of αSMA⁺ vessels in mLVRF-treated muscles, indicating increased neovascular maturation (**Figure 3B**). This was further supported by cross-sectional analysis of triceps muscles (**Figure 3C**), which showed a significant increase in vascular density (αSMA⁺/mm^2^) following mLVRF treatment (**Figure 3D**). Together, these findings demonstrate that mLVRF administration significantly improves microvascular architecture in *mdx* muscle.

### The combined LVRF+ASO treatment improves exon skipping efficacy and dystrophin restoration

To investigate whether improved vascularization enhances ASO treatment efficacy, we quantified ASO levels in muscle tissues. As expected, combining mLVRF with ASOs resulted in a 3.9-fold increase in ASO accumulation compared to ASO treatment alone (**Figure 4A**). This enhanced ASO delivery resulted in an 80% increase in exon skipping efficiency, as quantified by qPCR (**Figure 4B**), and a ∼50% increase in dystrophin restoration, as measured by western blot (**Figures 4C-D**). On average, dystrophin expression reached ∼ 10% in TA, gastrocnemius (GAS), quadriceps (QUAD), triceps (TRI) and diaphragm (DIA) muscles following ASO treatment alone, compared to ∼ 17% in the combined mLVRF+ASO group. Immunostaining confirmed correct sarcolemmal localization of dystrophin in TA, GAS, and DIA muscles (**Figure 4E**). Quantitative analysis of GAS sections further revealed a significant 15% increase in dystrophin-positive fibers in the combined treatment group compared to ASO alone (**Figure 4F**), reinforcing the beneficial impact of mLVRF co-administration. To evaluate the effect on muscle integrity, we measured serum Myomesin-3, a biomarker of muscle damage released into the circulation upon sarcolemmal disruption (**Figure 4G**). Serum Myomesin-3 levels were reduced by 46% with mLVRF alone, 59% with ASO alone, and up to 74% with the combined treatment, indicating a synergistic effect on sarcolemmal stabilization and muscle protection.

**Figure 4:**
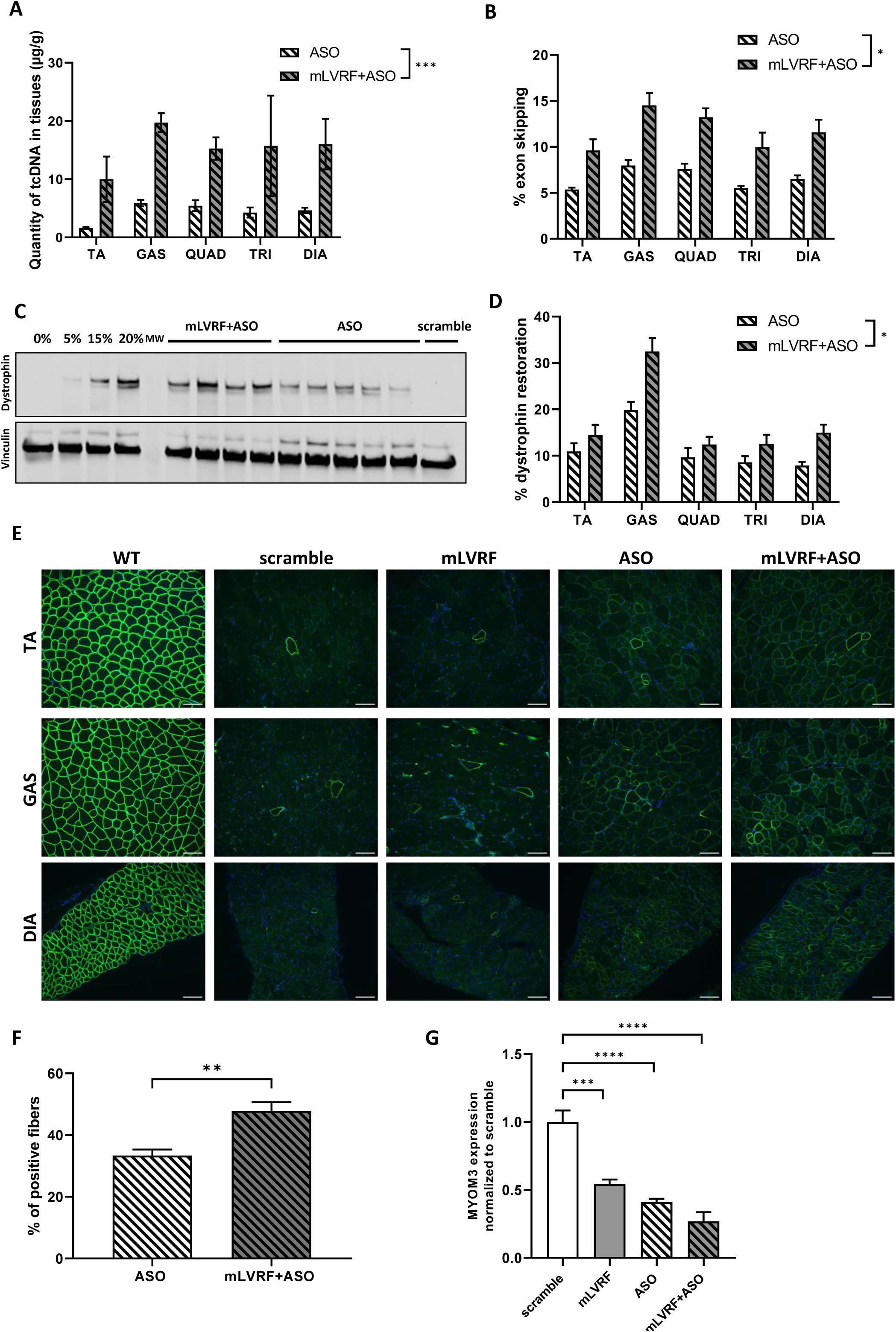
Combined treatment mLVRF+ASO improves dystrophin restoration. **(A)** Quantification of ASO in TA (tibialis anterior), GAS (gastrocnemius), QUAD (quadriceps), TRI (triceps) and DIA (diaphragm) after 12 weeks of ASO treatment. (***p<0.001 analyzed by two-way ANOVA). **(B)** Quantification by qPCR of exon 23 skipping in different muscles. (*p<0.05 analyzed by two-way ANOVA). **(C)** Representative WB from diaphragm protein lysates. A standard curve of 0%, 5%, 15% and 20% was made from pooled lysates from C57Bl10 (WT) and *mdx* for each tissue. **(D)** Quantification of dystrophin expression by western-blot (WB). (*p<0.05 analyzed by two-way ANOVA). **(E)** Dystrophin immunostaining on cross-sections of muscles from WT, scramble, mLVRF, ASO and mLVRF+ASO treated mice. Scale bar, 100µm. **(F)** Proportion of dystrophin positive fibers in GAS muscles. (**p<0.01, analyzed by t-test). **(G)** Western blot quantification of myomesin-3 (MYOM3) in serum after normalization to scramble group. Myomesin 3 is undetectable in the serum of WT mice. (***p<0.001; ****p<0.0001, analyzed by one-way ANOVA). Results are expressed as the mean ± SEM (n=5 in WT, scramble, mLVRF and ASO groups, n=4 in mLVRF+ASO group).

### Functional benefit and histopathology improvement

To assess the therapeutic benefit of the co-treatment, a series of functional tests were performed on treated mice. Endurance capacity was evaluated using a treadmill exhaustion test. As expected, the distance run by control *mdx* mice (scramble group) was significantly lower compared to WT mice (**Figure 5A**). ASO treatment alone led to a modest, non-significant improvement in endurance, whereas a statistically significant increase was observed only in the mLVRF+ASO group compared to scramble controls. A similar pattern was noted for maximum running speed, with significant improvement observed exclusively in the combined treatment group (**Figure 5B**).

**Figure 5:**
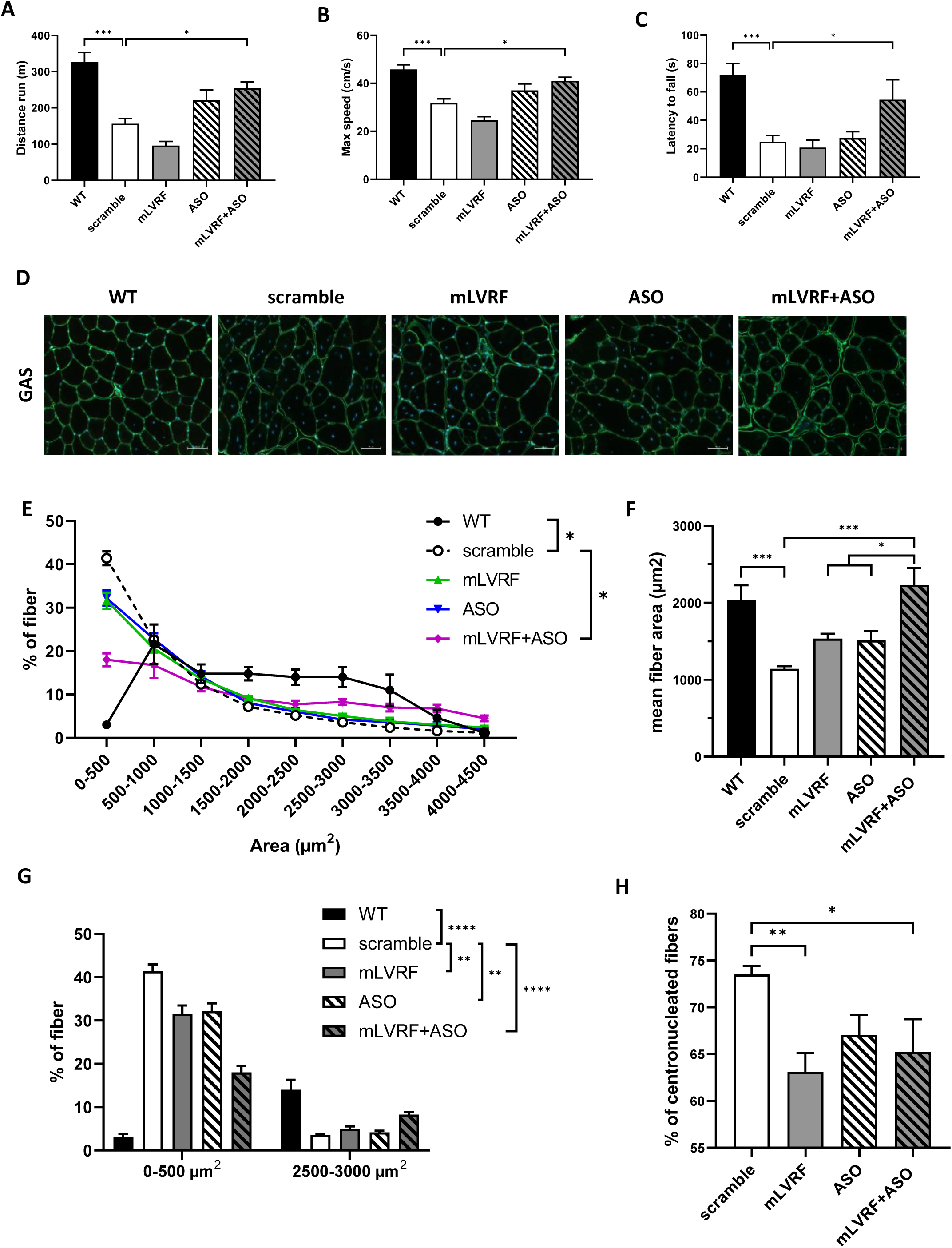
Impact of the mLVRF+ASO treatment on muscle function and histopathology. **(A)** Maximal distance run in meters on a treadmill. (*p<0.05; ***p<0.001 analyzed by one-way ANOVA). **(B)** Maximal speed reached by animals during the treadmill test. (*p<0.05; ***p<0.001 analyzed by one-way ANOVA). **(C)** Latency for each animal to fall from the inverted grid. (*p<0.05; ***p<0.001 analyzed by one-way ANOVA). **(D)** Laminin immunostaining of GAS muscles from WT, scramble, mLVRF, ASO and mLVRF+20-E treated mice. Scale bar, 100µm. **(E)** Representation of the percentage of fibers as a function of area including multiple intervals. Fiber sizes were quantified from the laminin staining. (*p<0.05, analyzed by two-way ANOVA). **(F)** Mean area of TA fiber size in the different groups. (*p<0.05, ***p<0.001, analyzed by one-way ANOVA). **(G)** Focus on the percentage of fibers in GAS muscles with an area between 0 and 500 µm^2^ or 2500 to 3000µm^2^. (**p<0.01; ****p<0.0001, analyzed by two-way ANOVA). **(H)** Percentage of centronucleated fibers in the different groups. WT mice are not presented because of the absence of centronucleated fibers. Results are expressed as the mean ± SEM (n=5 in WT, scramble, mLVRF and ASO groups, n=4 in mLVRF+ASO group).

To further assess functional benefit, mice were subjected to the inverted grid test, which measures global limb strength over a maximum period of 120 seconds. WT mice remained on the grid for an average of 72 seconds, while *mdx* scramble mice fell after only 24 seconds on average (**Figure 5C**). Among treated groups, only the combined mLVRF+ASO group showed a statistically significant improvement in latency to fall compared to scramble controls. Altogether, these results demonstrate that the combination of AAV-mLVRF and ASO treatment significantly improves both muscle endurance and strength in *mdx* mice, suggesting a clear therapeutic benefit of this combined therapeutic strategy.

We next evaluated the impact of mLVRF treatment on muscle histopathology (**Figure 5D**). As previously described, the fiber size distribution in *mdx* muscles differs markedly from that of WT muscles (**Figure 5E**). *Mdx* muscles display an increased proportion of small fibers (<500 µm²) and a reduced number of large fibers, reflecting excessive regeneration. Muscle fiber size distribution was significantly improved in mice treated with the combined mLVRF+ASO therapy compared to the scramble group. Similarly, the overall mean fiber area, typically reduced in *mdx* mice, was significantly increased only with the combined mLVRF+ASO treatment (**Figure 5F**), underscoring its beneficial effect on muscle fiber size. Both mLVRF and ASO monotherapies significantly reduced the proportion of small fibers and increased the number of larger fibers (**Figure 5G**). However, the combined treatment had a more pronounced effect by reducing small fibers to 18%, compared to 41% in the scramble group but also compared to the ASO alone group (p<0.0001 analyzed by two-way ANOVA).

Conversely, the proportion of larger fibers increased to 9% (compared to 3% in the scramble group) resulting in a size distribution more closely resembling that of WT muscles (**Figure 5G**). This histological improvement was further supported by a significant reduction in the percentage of centronucleated fibers, indicative of strong regeneration, in the mLVRF monotherapy and combined treatment groups (**Figure 5H**).

### Impact of the combined treatment on fibrosis and dystrophic biomarkers

A hallmark of fibrotic remodeling in *mdx* skeletal muscles is excessive collagen and other extracellular matrix deposition. To assess fibrosis, Sirius Red staining was performed on GAS cross-sections to quantify collagen content across treatment groups (**Figure 6A**). As expected, collagen content was markedly higher in scramble-treated *mdx* mice compared to WT controls. (**Figure 6B**). Both ASO monotherapy and the combined mLVRF+ASO treatment significantly reduced collagen deposition compared to scramble-treated controls. To further confirm these observations, expression levels of key fibrotic markers, including collagen I (Col1a1) and connective tissue growth factor (CTGF), were quantified by qPCR. A significant decrease in both markers was observed in the ASO and mLVRF+ASO groups compared to scramble controls (**Figure 6C**), supporting the anti-fibrotic effect of the treatments. In parallel, we evaluated the expression of several dystrophic biomarkers by qPCR. Expression of *Pax7*, a transcription factor essential for satellite cell maintenance and muscle regeneration, was significantly upregulated in *mdx* muscles compared to WT (**Figure 6D**). This overexpression reflects the chronic activation and expansion of the satellite cell pool in response to ongoing muscle damage. Although Pax7 is a hallmark of quiescent cells, its sustained expression in dystrophic conditions marks both activated and self-renewing satellite cells, consistent with a persistent regenerative attempt.^26^ Only the combined treatment with mLVRF and ASO led to a significant reduction in *Pax7* expression, suggesting a decreased regenerative burden and improved muscle homeostasis. Similarly, mRNA levels of *Troponin T*, encoding a sarcomeric protein elevated during muscle injury, were increased in *mdx* muscles, with significant normalization observed only following the combined mLVRF+ASO treatment (**Figure 6E**). We also investigated the expression of *PGC-1α*, a key regulator of mitochondrial biogenesis and oxidative metabolism. As expected, *PGC-1α* expression was significantly reduced in scramble-treated *mdx* mice compared to WT. Notably, only the combined treatment significantly restored *PGC-1α* levels (**Figure 6F**), indicating a potential improvement in the metabolic status of the muscle. Collectively, these findings underscore the beneficial effects of the combined mLVRF+ASO treatment on key histopathological features of dystrophic muscle, including fibrosis, regeneration, muscle damage, and metabolic alterations

**Figure 6:**
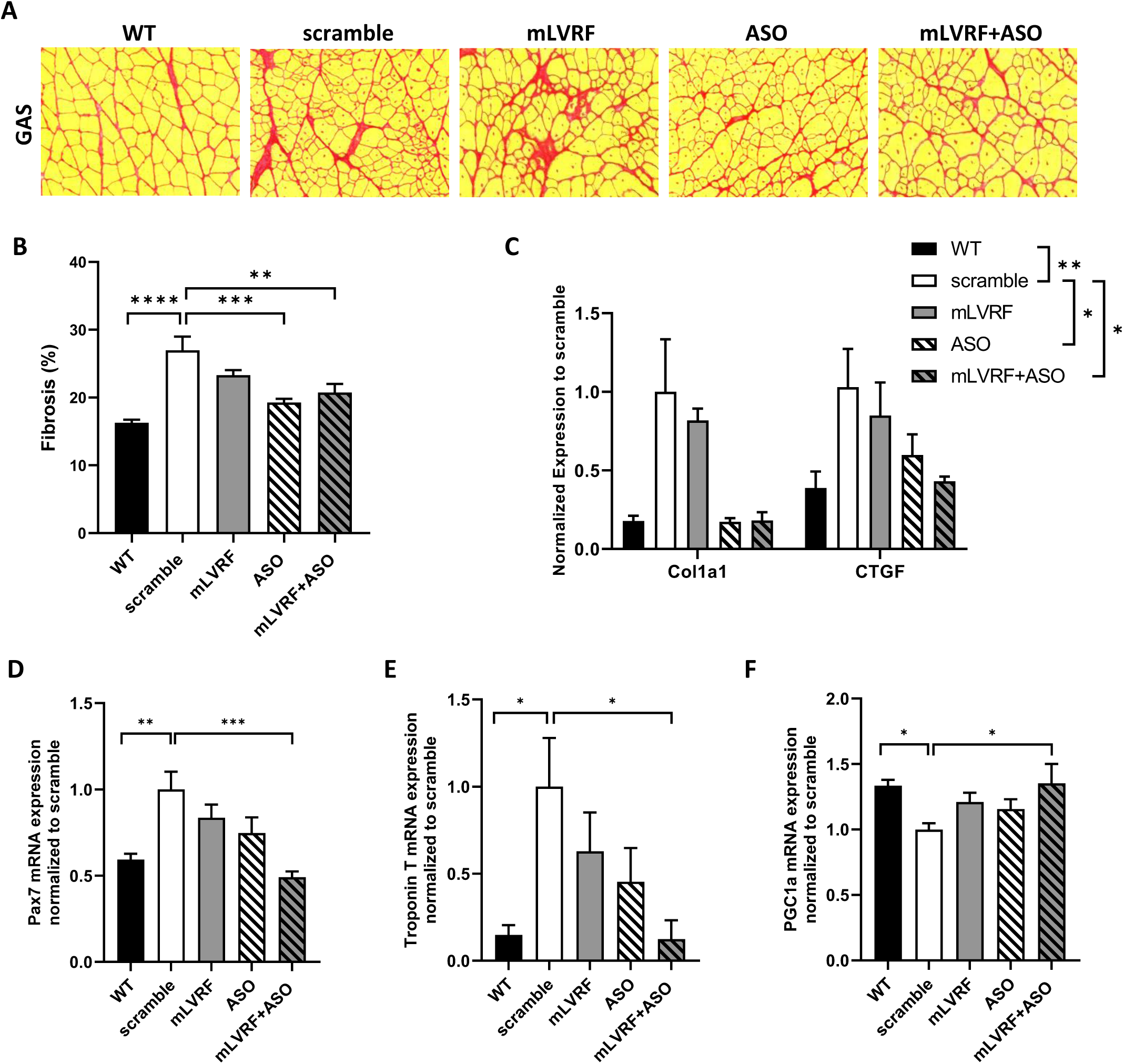
Impact of the combined treatment on fibrosis and dystrophic biomarkers. **(A)** Sirius red staining of GAS muscle from sections of WT, scramble, mLVRF, ASO and mLVRF+20-E treated mice. **(B)** Percentage of fibrosis found in different groups measured from Sirius red staining (**p<0.01; ***p<0.001; ****p<0.0001, analyzed by one-way ANOVA). (**C)** Quantification collagen 1 alpha1 subunit (Col1a1) and connective tissue growth factor (CTGF) expression by qPCR. (*p<0.05; **p<0.01, analyzed by two-way ANOVA). Quantification of the expression of Pax7 **(D)**, Troponin T **(E)** and PGC1a **(F)** in GAS muscle by qPCR. (*p<0.05; **p<0.01; ***p<0.001, analyzed by one-way ANOVA). Results are expressed as the mean ± SEM (n=5 in WT, scramble, mLVRF and ASO groups, n=4 in mLVRF+ASO group).

### Serum biochemistry

Finally, to evaluate potential treatment-related toxicity, serum biochemical markers were assessed in *mdx* mice across all groups (i.e AAV-mLVRF alone, ASO alone or the combined mLVRF+ASO). No significant changes were detected in serum urea, albumin, or creatinine (**Figure 7**), indicating preserved renal and hepatic function. Conversely, consistent with the overall therapeutic benefit, all treatments significantly reduced at least one marker of liver stress (alkaline phosphatase (ALP) and the transaminases ALT and AST) suggesting a potential systemic improvement associated with the therapy.

**Figure 7:**
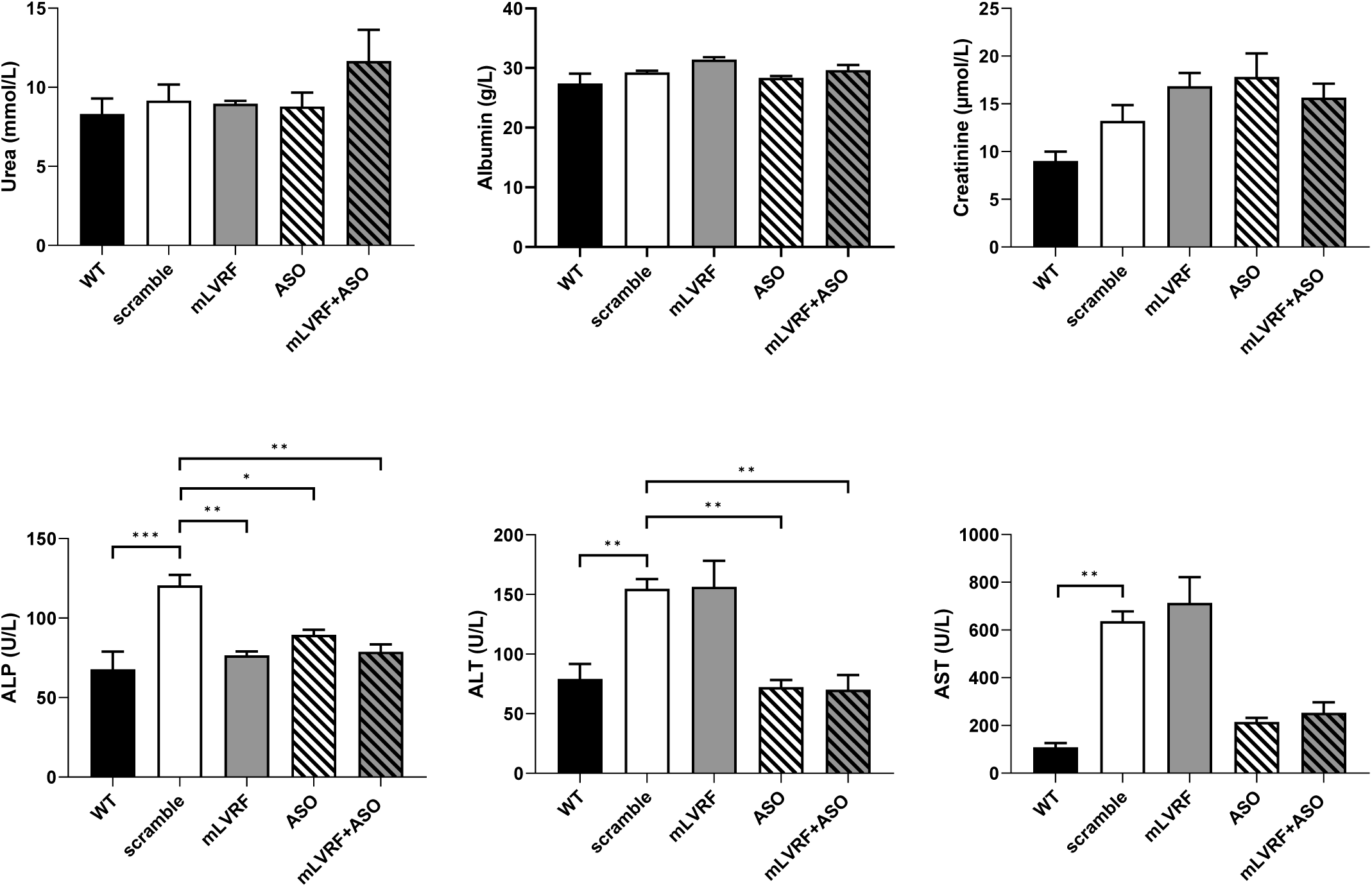
Serum biochemistry. Serum quantification of renal and liver toxicity markers including urea, albumin, creatinine, alkaline phosphatase (ALP), alanine transaminase (ALT), aspartate aminotransferase (AST). (*p<0.05; **p<0.01; ***p<0.001 analyzed by one-way ANOVA). Results are expressed as the mean ± SEM (n=5 in WT, scramble, mLVRF and ASO groups, n=4 in mLVRF+ASO group).

## DISCUSSION

Despite significant advances in ASO-based therapies for DMD, the clinical efficacy of these approaches remains limited by several factors, including inefficient delivery to skeletal muscles and heterogeneous biodistribution. Among the currently approved ASO therapies, eteplirsen (Exondys 51), golodirsen (Vyondys 53), viltolarsen (Viltepso), and casimersen (Amondys 45) have demonstrated modest increases in dystrophin restoration and functional outcomes, particularly in ambulatory patients. To be clinically meaningful, these therapies rely on efficient ASO delivery to muscles from the vascular compartment after ASO intravenous infusion, which is challenged by the progressive muscle degeneration and poor vascularization characteristic of DMD. Many international efforts are currently focusing on improving the targeted delivery of ASO to muscle tissues, notably conjugating the ASO to targeting moieties such as peptides, lipid or antibodies. Several biotechnology companies have initiated programs to enhance ASO delivery to muscle by developing proprietary conjugation platforms. Among these, peptide-conjugated PMOs (PPMOs) have shown promise for improving cellular uptake and endosomal escape. For example, Sarepta’s PPMO vesleteplirsen (SRP-5051) demonstrated enhanced exon-skipping and dystrophin production in Phase 2 trials compared to eteplirsen (NCT04004065); however, Sarepta announced in late 2024 that it would discontinue the program due to long-term safety and tolerability considerations. Similarly, PepGen’s PGN-EDO51 was designed to balance efficacy and safety, and showed encouraging initial results in healthy volunteers, but the company recently decided to end its development for DMD following limited dystrophin restoration and some kidney toxicity observed in the Phase 2 trial (NCT06079736). In parallel, the development of antibody–oligonucleotide conjugates has gained significant traction and may offer promising opportunities to more efficiently target muscle tissues. For example, Dyne Therapeutics is advancing DYNE-251, an ASO conjugated to a transferrin receptor–binding Fab fragment, which is being evaluated in the global Phase 1/2 DELIVER trial in individuals with DMD amenable to exon 51 skipping. Notably, Dyne recently announced encouraging long-term data demonstrating unprecedented and sustained functional improvement at the selected registrational dose, underscoring the potential of antibody-based targeting strategies.^27^ Avidity Biosciences has similarly advanced an antibody-oligonucleotide conjugate (AOC-1044, also known as delpacibart zotadirsen) targeting DMD exon 44 in the Phase 1/2 EXPLORE44 trial.^28,29^ Despite these promising advances in ASO delivery, effective targeting of dystrophic muscles remains constrained by the inherently compromised microvasculature characteristic of DMD. In this study, we demonstrate that improving the muscle microenvironment through targeted angiogenesis significantly enhances the efficacy of ASO therapy. We show that AAV-mediated delivery of the pro-angiogenic LVRF increases microvascular density, restores capillary-to-fiber ratios closer to wild-type levels, and facilitates ASO biodistribution and uptake across multiple muscle groups. This improvement translates into enhanced exon skipping efficiency and dystrophin restoration, ultimately resulting in better muscle histopathology and functional performance. Notably, we observed improvements in muscle fiber size distribution, a reduction in the proportion of centronucleated fibers, decreased fibrosis, and enhanced performance in functional assays such as treadmill and inverted grid tests.

It is worth noting that while the combined pro-angiogenic and ASO treatment resulted in a substantial 3.8-fold increase in ASO biodistribution compared to ASO alone, this translated into a more modest 1.8-fold improvement in exon skipping efficiency. This discrepancy suggests that although larger amounts of ASO successfully reach the muscle tissue through improved vascularization, other well-known barriers, such as limited productive cellular uptake and sequestration within the endo-lysosomal compartment, still constrain their full therapeutic effect. These results highlight that further strategies targeting intracellular delivery and endosomal escape could work synergistically with enhanced vascularization to achieve even greater levels of dystrophin restoration.

Importantly, LVRF treatment appeared well tolerated in *mdx* mice, alone or in combination with ASO, contrasting with VEGFA treatment, which was not, even injected at lower dose, and thus prevented further side to side comparison. Moreover, the comparative analyses performed in our dose-finding study reveal that LVRF has a more favorable impact on muscle vasculature, without the deleterious side effects associated with excessive VEGFA signaling. These findings underscore the potential of LVRF as a safer and more efficient angiogenic modulator in dystrophic muscles. An additional advantage of LVRF is its ability to engage both VEGFA and VEGFC pathways, thereby promoting not only blood vessel formation but also lymphangiogenesis. Lymphatic vessels play a crucial role in interstitial fluid clearance and immune cell trafficking, but emerging studies suggest they may also facilitate the recycling and systemic redistribution of ASOs.^30^ Enhancing lymphatic capacity in dystrophic muscle might therefore not only improve fluid homeostasis and reduce inflammation but also support ASO bioavailability and retention in target tissues. Taken together, dual angiogenic/lymphangiogenic modulation by LVRF could significantly augment both tissue remodeling and therapeutic distribution in neuromuscular disease contexts. Given the intrinsic vascular deficiency in DMD, strategies aimed at restoring a healthy muscle vasculature should be considered as valuable adjuncts to current and future therapeutic interventions. The benefits observed with ASO delivery could likely extend to other systemic gene therapies such as AAV-based microdystrophin delivery, which also depends on effective intravascular distribution and transduction. The recent accelerated approval of *Elevidys* marks a significant milestone in the treatment of DMD. However, the high vector doses required for therapeutic efficacy remain a major concern, particularly in light of the recent deaths of two patients enrolled in clinical trials. These events highlight the urgent need for strategies that can reduce the required AAV doses. By improving muscle microvasculature, the approach described here could enhance vector delivery and distribution, thereby contributing to dose reduction and improved safety in systemic AAV-based therapies.

While our proof-of-concept study utilized AAV vectors to deliver the mLVRF transgene, we acknowledge the limitations associated with sustained overexpression and potential physiological perturbations, especially considering possible future combinations with other AAV-based gene therapies such as AAV-microdystrophin. Therefore, future research should explore alternative approaches to induce mLVRF activity without relying on viral overexpression. One promising avenue could involve splice-switching strategies that favor the endogenous production of the LVRF isoform through modulation of VEGFA alternative splicing, potentially using ASO or small molecules. Moreover, beyond VEGFA, other angiogenesis-promoting pathways could be targeted to avoid the risks associated with VEGFA overexpression, such as vascular permeability or aberrant angiogenesis. For example, modulating Angiopoietin-Tie2 signaling or using prolyl hydroxylase inhibitors to stabilize HIF pathway could represent alternative or complementary strategies to achieve vascular normalization in dystrophic muscles.^31,32^

In conclusion, our findings provide compelling evidence that correcting the vascular niche significantly enhances ASOs therapeutic efficacy in DMD. Notably, even modest increases in exon skipping and dystrophin restoration can be sufficient to convert a marginally effective approach into a clinically meaningful treatment, especially given the limited efficacy of currently approved PMO-ASOs. This combinatorial approach hold promise not only for ASO-based therapy but also for other systemic treatments. Future efforts should focus on refining angiogenesis-targeted interventions to maintain physiological balance, potentially through precise isoform regulation or modulation of alternative pathways.

## MATERIALS & METHODS

### Antisense Oligonucleotides and Animal Experiments

All procedures involving animals were conducted in accordance with national and European regulations and the ARRIVE guidelines and were approved by the French Ministry of Research (APAFiS #6518). *Mdx* mice (C57BL/10ScSc-Dmdmdx/J) were bred and housed under standard conditions at the 2Care platform (Université de Versailles Saint-Quentin).

TcDNA antisense oligonucleotides (ASOs) targeting the donor splice site of exon 23 of the murine *Dmd* pre-mRNA (5′-CCTCGGCTTACCT-3′) were synthesized by SQY Therapeutics (Montigny-le-Bretonneux, France). A palmitic acid moiety was conjugated to the 5′ end of full phosphodiester tcDNA via a C6-amino linker and a terminal phosphorothioate bond as previously described.^12^

Seven-to ten-week-old male *mdx* mice received a single intravenous injection of AAV9 encoding either VEGF isoform A164 (1E+13 vg/kg) or 234NF (3E+13 vg/kg) four weeks prior to ASO administration. Mice were subsequently treated intravenously with tcDNA-ASO (50 mg/kg/week) for 12 weeks. Control groups included *mdx* mice injected with AAV9-scramble followed by saline, and age-matched C57BL/10 wild-type mice. One week after the final ASO dose, mice were euthanized by cervical dislocation. Tissues were collected, snap-frozen in liquid nitrogen-cooled isopentane, and stored at −80°C.

### Functional Analysis

#### Treadmill Test

Mice were acclimated to the test room for 30 minutes prior to each session. For three consecutive days before testing, mice were exposed to a non-moving treadmill for 30 seconds followed by a 5-minute warm-up at 20 cm/s. On the test day, a 3-minute warm-up at 5 cm/s was followed by incremental speed increases of 1 cm/s every 30 seconds until exhaustion. Exhaustion was defined as the point when mice failed to resume running within 20 seconds despite gentle prodding. Total running distance was recorded for each animal.

#### Inverted Grid Test

Following a 30-minute acclimation period, mice were placed on a metal grid, which was then inverted to assess grip strength and endurance. Each mouse underwent three trials, with 300 seconds of rest between trials. The test was capped at 120 seconds per trial, and latency to fall was recorded.

### ASO Quantification by Fluorescent Hybridization Assay

Tissues were homogenized using the Precellys system (Bertin Instruments, Montigny-le-Bretonneux, France) in lysis buffer (100 mM Tris-HCl, pH 8.5; 200 mM NaCl; 5 mM EDTA; 0.2% SDS) containing 2 mg/ml proteinase K (Invitrogen, Carlsbad, USA) at a ratio of 50 mg tissue per ml of buffer. Homogenates were incubated overnight at 55°C, then centrifuged at 7,000 rpm. Supernatants were collected for ASO analysis. ASO concentrations were measured using a molecular beacon-based fluorescent hybridization assay, as previously described. Briefly, 10 µl of each lysate was incubated in black non-binding 96-well plates (Fisher Scientific, Waltham, USA) with a 5′-Cy3-labeled DNA probe complementary to the ASO and a 3′ HBQ quencher. Reactions were brought to 100 µl with PBS, and fluorescence was measured using a FluoStar Omega plate reader (BMG Labtech, Ortenberg, Germany; Ex 544 nm / Em 590 nm). ASO levels were determined by interpolation from a standard curve generated with known concentrations of tcDNA diluted in lysates from untreated tissues.

### RNA Analysis

Total RNA was extracted from snap-frozen muscle tissues using TRIzol reagent (ThermoFisher Scientific, Waltham, USA), following the manufacturer’s instructions. Exon 23 skipping was quantified using the QX200 Droplet Digital PCR (ddPCR) system (Bio-Rad, Hercules, USA). One microgram of RNA was reverse-transcribed using the LunaScript RT SuperMix kit (New England Biolabs, Ipswich, USA), and 100 ng of cDNA were used per ddPCR reaction. Amplification was performed using the ddPCR Supermix for probes (no dUTP) with specific probes targeting the exon 23–24 and exon 22–24 junctions. Droplet generation was conducted using the QX200 Droplet Generator, and PCR was carried out as per the manufacturer’s protocol. Droplets were analyzed using the QX200 Droplet Reader. Quantitative PCR (qPCR) was also performed on diluted cDNA (1:10) using the iTaq Universal SYBR Green Supermix (Bio-Rad, Hercules, USA) to assess the expression of *VEGF*, *LVRF*, and genes associated with regeneration, fibrosis, angiogenesis, and metabolism. Primer sequences are provided in **Table S1**.

### Western Blot Analysis

Frozen muscle tissues were cryosectioned and homogenized using the Precellys 24 system (Bertin Instruments, Montigny-le-Bretonneux, France) in RIPA buffer supplemented with 5% SDS and protease inhibitors. Lysates were denatured, centrifuged, and supernatants were collected. Total protein concentration was determined using the BCA Protein Assay Kit (ThermoFisher Scientific, Waltham, USA). For dystrophin detection, 25 µg of protein were resolved on NuPAGE 3–8% Tris-Acetate Protein Gels (Invitrogen, Carlsbad, USA), then transferred and blocked using EveryBlot Blocking Buffer (Bio-Rad, Hercules, USA). Membranes were incubated with NCL-DYS1 monoclonal anti-dystrophin antibody (1:1000, Novocastra, Newcastle upon tyne, UK) and hVin-1 anti-vinculin antibody (1:25,000, Sigma, St. Louis, USA), followed by IRDye 800CW goat anti-mouse IgG secondary antibody (1:2000, Li-Cor, Lincoln, USA). Visualization was performed using the Odyssey CLx Imaging System (Li-Cor, Lincoln, USA), and quantification was carried out using Empiria Studio software, referencing a standard curve generated from pooled lysates of C57BL/10 and mdx control muscles.

To detect circulating myomesin-3, mouse serum samples were diluted 1:20 and separated on 3–8% Criterion™ XT Tris-Acetate Protein Gels (Bio-Rad, Hercules, USA). Western blotting was performed using the iBind™ Flex Western Device (Fisher Scientific, Waltham, USA). Membranes were incubated with a rabbit polyclonal anti-MYOM3 antibody (1:1000, Proteintech, Rosemont, USA), followed by IRDye 800CW goat anti-rabbit IgG secondary antibody (1:2000, Li-Cor, Lincoln, USA). Bands were visualized using the Odyssey Imaging System (Li-Cor, Lincoln, USA). Signal intensity was normalized to total protein staining using Revert 700 Total Protein Stain (Li-Cor, Lincoln, USA) and further normalized to PBS-treated controls using Image Studio software.

### VEGF Quantification by ELISA

Tissue VEGF protein levels were quantified using the Quantikine® Mouse VEGF ELISA kit (MMV00, R&D Systems, Minneapolis, MN, USA), according to the manufacturer’s instructions. Samples were homogenized in lysis buffer containing 50 mM Tris-HCl (pH 7.4), 250 mM NaCl, 5 mM EDTA, 1% Triton X-100, and a protease inhibitor cocktail. Following centrifugation, the supernatants were collected, and total protein concentrations were determined using the BCA Protein Assay Kit (Thermo Fisher Scientific, Waltham, USA). VEGF levels were normalized to total protein content and expressed as pg/mg of protein.

### Immunohistochemistry

Cryosections (10 µm thick) were collected every 120 µm and stained for dystrophin, laminin, CD31, and α-SMA. Sections were incubated with the following primary antibodies: rabbit polyclonal anti-dystrophin (1:500, ThermoScientific, Waltham, USA, RB-9024-P), rabbit polyclonal anti-laminin (1:200, Sigma, St.Louis, USA, L9393-2ML), and rat anti-CD31 (1:1000, BD Pharmingen, San Jose, USA, 550274). Corresponding secondary antibodies included goat anti-rabbit IgG (H+L) F(ab’)₂ fragment Alexa Fluor® 488 (1:500–1:1000) and goat anti-rat IgG (H+L) Alexa Fluor® 555 (1:1000). All images were captured at equivalent exposure times and analyzed using ImageJ and Imaris software.

For Picro Sirius Red staining, 10 µm frozen sections were air-dried for 1 hour, treated with xylene for 10 minutes to reduce red fiber overstaining, and rehydrated through a graded ethanol series (100%, 80%, 40%). Sections were incubated in distilled water for 1 minute, stained with Picro Sirius Red solution (Abcam, Cambridge, UK, ab246832) for 1 hour, and rinsed twice with 0.1N HCl. Dehydration was performed using an ethanol gradient (70%, 80%, 100%), followed by clearing in xylene and mounting with Vectamount® permanent mounting medium (Vector Laboratories, Newark, USA). Images were acquired using a Leica DFC7000 T camera and Leica DM IL microscope (Leica Microsystems, Wetzlar, Germany), and analyzed with ImageJ (v1.54d).

### Muscle Dissociation and Flow Cytometry Analysis

Muscle tissues were harvested from mice and immediately placed on ice in a Petri dish containing HBSS supplemented with 1% penicillin-streptomycin (HBSS+). Muscles were finely minced (<2 mm pieces) and transferred into 15 mL conical tubes containing 8 mL of HBSS+. Samples were centrifuged at 1300 rpm for 1 min at 4 °C. The supernatant was discarded, and 4 mL of digestion buffer was added to the tissue pellet. Samples were incubated horizontally at 37 °C for 30 min. After a brief settling, the supernatant was collected and subjected to a second 30 min incubation at 37 °C under agitation. Digestion was stopped by adding HBSS+ up to a final volume of 15 mL. Cell suspensions were filtered sequentially through 100 µm and 70 µm cell strainers and centrifuged at 300 g for 5 min at 4 °C. The pellet was resuspended in 500 µL PBS, and 200 µL aliquots were set aside for viability controls. Viability staining was performed using the LIVE/DEAD™ Fixable Dead Cell Stain Kit (ThermoFisher Scientific, Waltham, USA) at a 1:2000 dilution. Samples were incubated for 3 min at room temperature, quenched with 5 mL of HBSS+, and centrifuged at 1300 rpm for 5 min at 4 °C. Cells were fixed in 4% paraformaldehyde for 15–20 min at room temperature, washed twice with HBSS+, and finally resuspended in 500 µL HBSS+ and stored at 4 °C overnight. A 10 µL aliquot of each tube was used to determine cell count. At least 50,000 cells per tube were used for compensation controls. Cells were blocked with 5% mouse serum (or FBS) for 5 min on ice, followed by incubation with fluorophore-conjugated antibodies diluted 1:200. The following antibodies were used: anti-CD31 (clone 390, PE-conjugated, Ozyme, Saint-Cyr-l’École, France, ref. TNB35-0451-U100) and anti-CD45 (clone 30-F11, FITC-conjugated, Thermo Fisher Scientific, Waltham, USA, ref. 15268509). Samples were incubated for 30 min at room temperature in the dark. After incubation, cells were washed with 3 mL HBSS+, centrifuged at 1300 rpm for 5 min at 4 °C, and resuspended in 500 µL of HBSS+ containing 2 mM EDTA. Samples were maintained on ice and immediately analyzed by flow cytometry.

### LEC Sprouting Assay

Human lymphatic endothelial cells (LECs) sprouting was assessed using a 3D fibrin matrix. Briefly, LECs were aggregated onto gelatin-coated microcarrier beads (250 beads per 2×10⁵ cells) by incubation overnight at 37 °C in Microvascular Endothelial Cell Growth Medium (PELOBiotech, Planegg/Martinsried, Germany). The following day, LEC-coated beads were embedded in a fibrin matrix composed of bovine fibrinogen and thrombin. After polymerization, human fibroblasts (FHN, 20,000 cells/well) were seeded on top of the polymerized gels in endothelial cell medium without VEGF and treatments with recombinant human LVRF, VEGF-A, or VEGF-C (200 ng/mL) were applied. After 6–7 days of incubation, samples were fixed with 4% paraformaldehyde for 1 h at room temperature, permeabilized with 0.5% Triton X-100 in PBS for 45 min, and stained overnight at 4 °C with Alexa Fluor 488– conjugated phalloidin. Sprouting was quantified using fluorescence microscopy as sprout-positive beads per well.

### Tissue clearing and immunolabelling

To visualize the vasculature, mice were intravenously injected with 200 μL of Lycopersicon Esculentum (Tomato) Lectin conjugated to DyLight649 (Eurobio, Les Ulis, France, DL-1178) 30 minutes prior to sacrifice. TA muscles were then immediately isolated and fixed in 4% paraformaldehyde (PFA) for 2 hours at 4°C under gentle rotation. Fixed tissues were washed three times in PBS for 10 minutes at 4°C under agitation. Muscles were longitudinally sliced into ∼1 mm-thick sections and directly processed for tissue clearing. Samples were incubated in RapiClear® 1.52 clearing reagent (SunJin Lab, Hwaseong-si, South Korea) for at least 12– 24 hours at room temperature in the dark. Imaging was performed using a CSU-W1 Spinning Disk confocal microscope (Nikon, Tokyo, Japan) with a 20× dry objective.

Clearing of the Extensor Digitorum Longus (EDL) muscle of mice was performed following the SUMIC protocol (Simple Ultrafast Multicolor Immunolabelling and Clearing) with some adaptations.^33^ Briefly, freshly dissected muscles were fixed in ice-cold paraformaldehyde/glutaraldehyde solution, then dehydrated through an increasing methanol gradient, bleached in a hydrogen peroxide solution, and rehydrated through a decreasing methanol gradient. After antigen retrieval and permeabilization, collagenase A-based matrix digestion was performed prior to the immunolabelling process, which included blocking, immunostaining, and washes. Immunolabelling was carried out using an anti-αSMA-CY3 antibody (Sigma, St.Louis, USA, C6198) at a 1:250 dilution for 2.5 days at 37°C. EDL muscles were embedded in 1% low melting agarose using a 1 ml syringe with a cut tip, dehydrated through an increasing isopropanol gradient, and finally cleared in ethyl cinnamate (ethyl 3-phenyl-2-propenoate; ECi). The cleared samples were imaged using a Zeiss Lightsheet 7 microscope equipped with dual-sided right/left light sheet illumination (5× objective, NA 0.1) and a detection objective (5×, NA 0.16) connected to an Axio Cam 701 camera (1920 × 1216 pixels). For image acquisition, the agarose-embedded cleared samples were manually affixed to the sample holder adapter using a drop of Superglue, then immersed in ECi within the sample chamber. Illumination was provided by light sheets at a wavelength of 561 nm, with laser intensity set to 10%. Three-dimensional image stacks were processed using Zen Lite 3.7 software to generate maximum intensity projections.

### Serum Analysis

Biochemical analysis of serum parameters—including alanine aminotransferase (ALT), aspartate aminotransferase (AST), alkaline phosphatase (ALP), creatinine, urea, and albumin— was conducted by the pathology laboratory at the Mary Lyon Centre (Medical Research Council, Harwell, Oxfordshire, UK).

### Viral Genome Quantification

Genomic DNA was isolated from tissue samples using the Gentra Puregene Tissue Kit (Qiagen, Hilden, Germany), in accordance with the manufacturer’s protocol. Quantification of viral genome copy number was carried out via TaqMan-based qPCR. The viral genome load was normalized to the number of transferrin receptor (TFRC1) gene copies to account for differences in genomic DNA input.

### Statistical Analysis

All *in vivo* data were analyzed using GraphPad Prism 8 (GraphPad Software, San Diego, CA, USA) and are presented as mean ± standard error of the mean (SEM). The number of animals per group is indicated as “n” in the figure legends. Group comparisons were made using one-way or two-way analysis of variance (ANOVA), with repeated-measures ANOVA applied where appropriate. To evaluate the overall effect of treatments across multiple tissues, two-way ANOVA was used, and the corresponding treatment *P*-value is reported in the figure legend. When data did not meet the assumptions of normality (as determined by the Shapiro–Wilk test), the non-parametric Kruskal–Wallis test was employed. Statistical significance was defined as follows: *p < 0.05, **p < 0.01, ***p < 0.001, ****p<0.0001.

### Data availability statement

The primary data for this study are available from the authors upon request.

## Acknowledgments

This work was supported by Institut National de la santé et la recherche médicale (INSERM), Université Paris-Saclay (France), Association Monégasque contre les myopathies (AMM), Paris Ile-de-France Region, and a PhD fellowship from Ministère de l’Enseignement Supérieur et de la Recherche (France) to M.B. We would like to thank the personnel of the platform 2CARE for taking care of the animals used in this work. We acknowledge the ImagoSeine core facility of the Institut Jacques Monod, member of the France BioImaging infrastructure (ANR-10-INBS-04) and GIS-IBiSA.

## Author contributions

Conceptualization and methodology, M.B., G.P. and A.G.; Investigation, M.B., M.D., C.G., X.P., O.LC., A.R., S.BA., and M.D.; Writing – Review & Editing, M.B., G.P. and A.G; Funding Acquisition, L.G. and A.G; Supervision A.G.

