## Supplemental material for "Improving angiogenesis ameliorates the efficacy of ASO-based exon-skipping for the treatment of Duchenne muscular dystrophy"

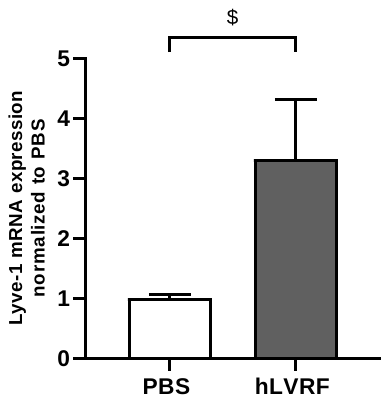

**A**

**C**

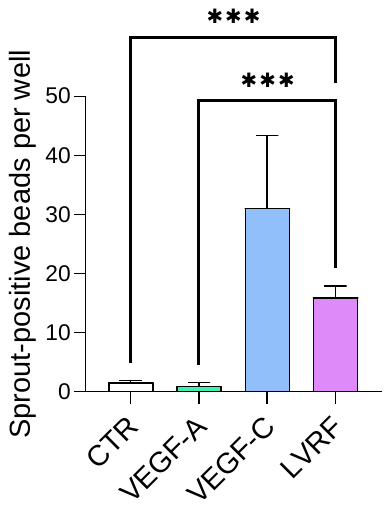

**B**

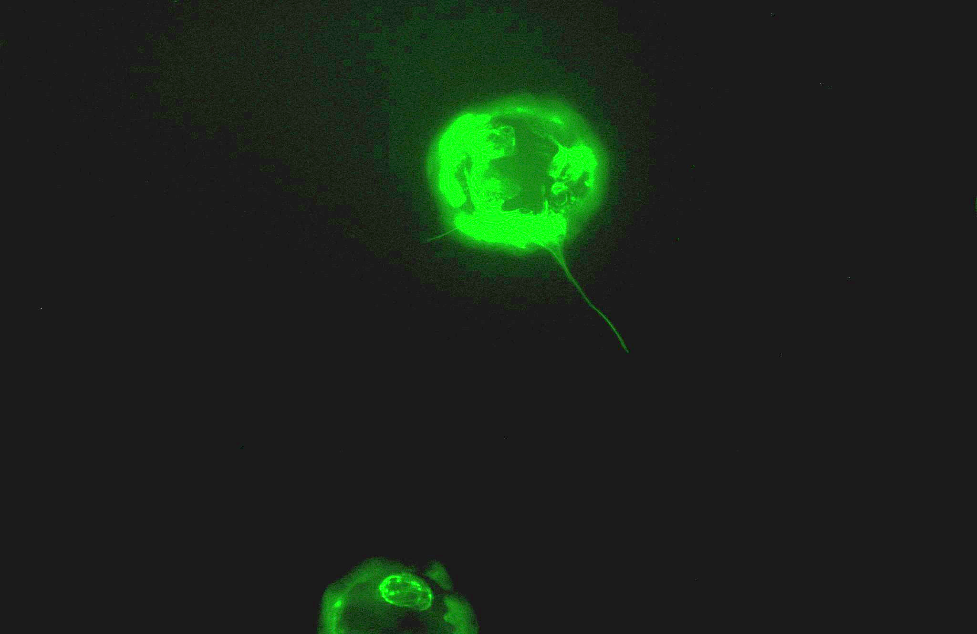

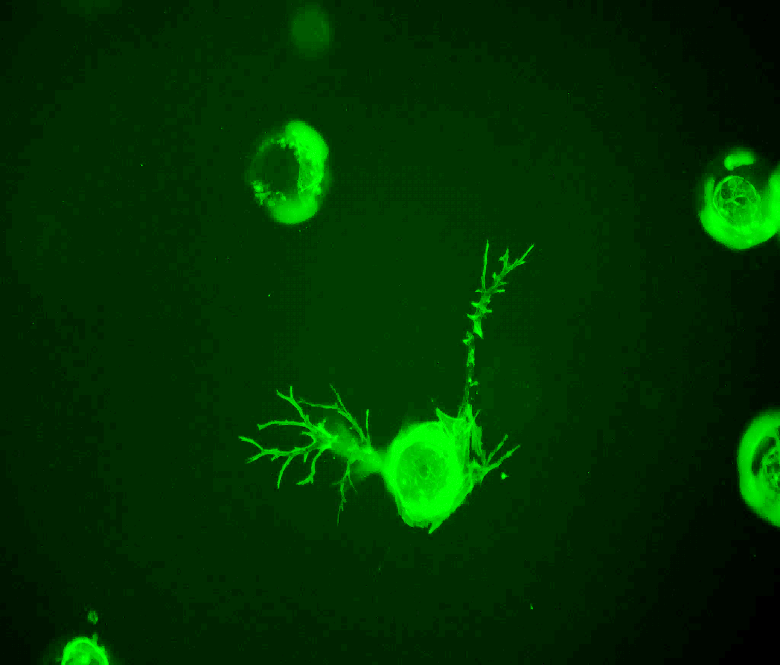

CTRL

LVRF

1

2

3

4

5

6

7

8

1

2

3

4

5

6

7

8

NF

VEGFA165

hLVRF

**D**

**Figure S1**: **Exon composition of hLVRF and pro-lymphangiogenic** activity. (A**)** Representation of the difference of exon composition between hLVRF and VEGFA165 isoforms. **(B)** Fluorescence microscopy of beads for control and LVRF-treated conditions. **(C)** Number of sprout-positive beads after stimulation with VEGF-A, VEGF-C or LVRF. n=3 per group. (***p<0.001, analyzed by one-way ANOVA). **(D)** Lyve-1 expression quantification by qPCR. n=4 per group. ($ p=0.0578, analyzed by t-test). Results are expressed as the mean ± SEM

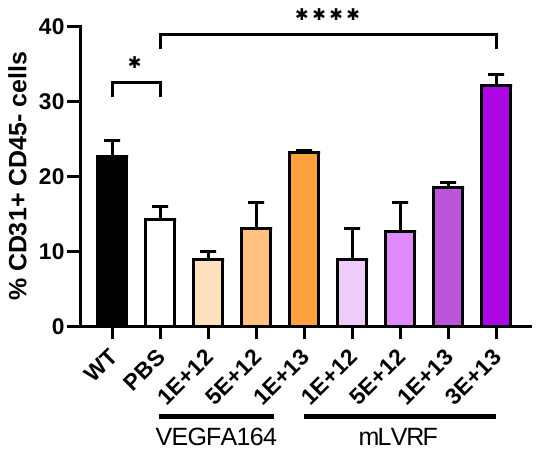

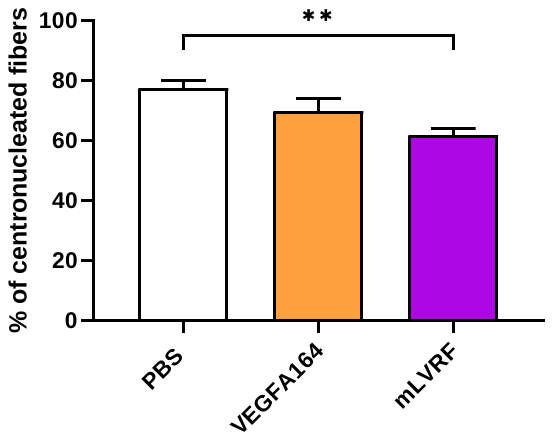

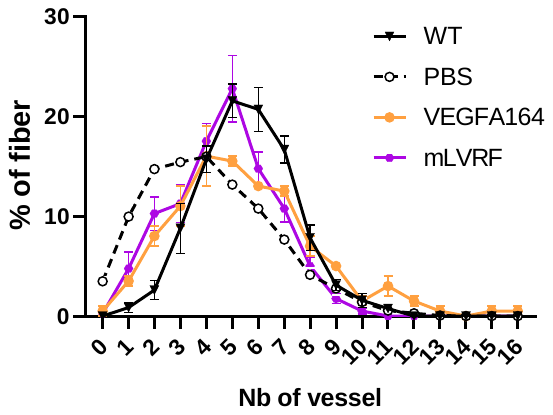

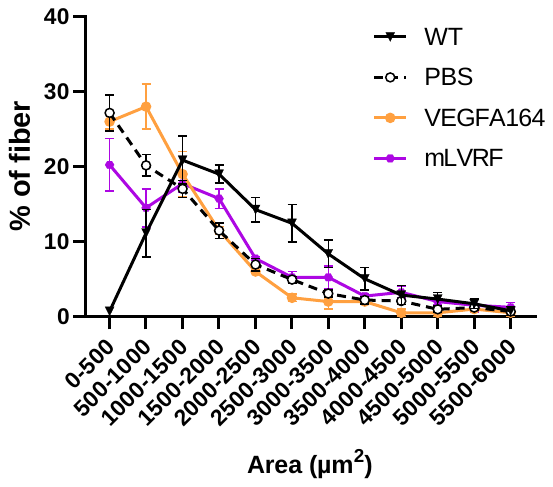

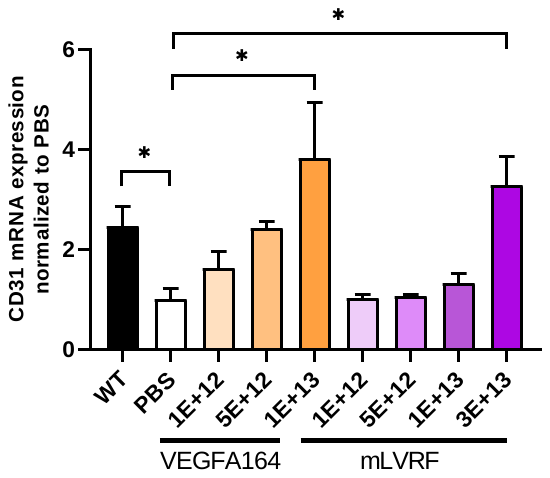

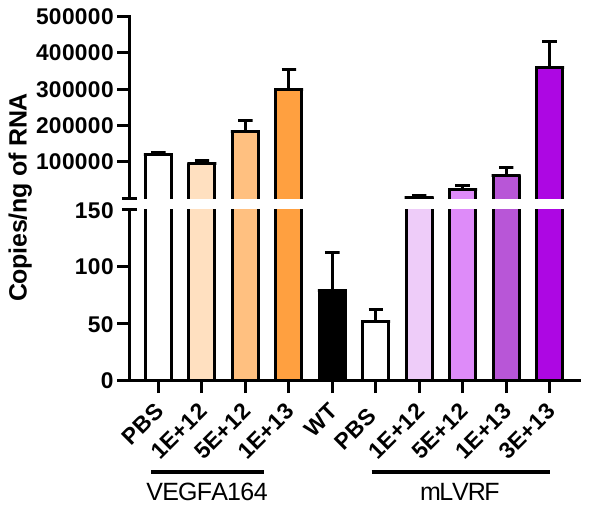

**A**

**B**

**C**

**D**

**E**

**F**

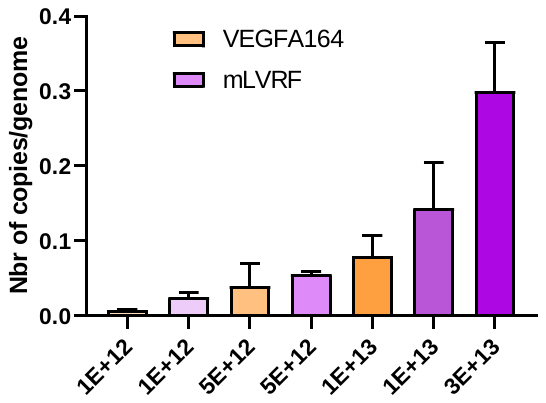

**G**

**Figure S2**: **mLVRF promotes angiogenesis and modulates muscle regeneration *in vivo*. (A)** Quantification by qPCR of the number of viral genomes normalized to the number of *TFRC1* copy in TA muscles. **(B)** Quantification by qPCR of the absolute number of VEGF A164 or mLVRF copies in TA. **(C)** qPCR quantification of CD31 expression after VEGF injections in TA lysates. (*p<0.05, analyzed by one-way ANOVA). **(D)** Flow cytometry quantification of endothelial cells in the gluteus muscle. (*p<0.05; ****p<0.0001, analyzed by one-way ANOVA). For the next figures (**E, F, G**), we only analysed the doses of 1E+13vg/kg for the VEGFA164 and 3E+13vg/kg for the mLVRF. (**E)** Distribution of the percentage of fibers as a function of the number of associated blood vessels. (**F)** Representation of the percentage of fibers as a function of area including several intervals. The data for angiogenesis and fiber size were obtained through laminin and CD31 co-staining of tibialis anterior (TA) cross-sections. (**G)** Percentage of centronucleated fibers in the different groups. WT mice are not presented because of the absence of centronucleated fibers. (**p<0.01, analyzed by one-way ANOVA). Results are expressed as the mean ± SEM (n=3 per group).

|  | **Target** | **Forward (5'** –> 3') | **Reverse (5' –> 3')** | **Probe (5' –> 3')** |
| --- | --- | --- | --- | --- |
| TAQMAN | DMD exon junction (23-24) | GAAACTTTCCTCCCAGTTGGT | CAGGCCATTCCTCTTTCAGG | GCCATCCATTTCTGTAAGGT |
|  | DMD exon junction (22-24) | CTGAATATGAAATAATGGAGGAGAGACTCG | CTTCAGCCATCCATTTCTGTAAGGT | TGTAATTTCC |
| SYBER | Collagen I | CATTGTGTATGCAGCTGACTTC | CGCAAAGAGTCTACATGTCTAGG |  |
|  | CTGF | TTGACAGGCTTGGCGATT | GTTACCAATGACAATACCTTCTGC |  |
|  | mLVRF | AGGCTCCCGTGGCCCTAAC | TCAACGGTGACGATGATGGCG |  |
|  | Troponin T | CCACCGCTTCCTCATCTTC | TCAGACTCCCTCTACAGCATC |  |
|  | CD31 | ACGCTGGTGCTCTATGCAAG | TCAGTTGCTGCCCATTCATCA |  |
|  | hLVRF | CCTTTGTTTTCCATTTCCCT | TCTGTCGATGGTGATGGTGTG |  |
|  | PGC1a | TCGCTCAATAGTCTTGTTCTCAA | AGAAGTCCCATACACAACCG |  |
|  | VEGFR3 | AGGTCAGAGAAAGCAGGTCT | CTGAAGGATGGCACTCGAAT |  |
|  | PAX7 | GAAGAAGTCCCAGCACAGC | GCTACCAGTACAGCCAGTATG |  |
|  | GAPDH | AATGGTGAAGGTCGGTGTG | GTGGAGTCATACTGGAACATGTAG |  |

**Supplementary table 1: Probes sequences used in taqman or sybergreen assays**
